# The cortical scene processing network emerges in infancy, prior to independent navigation experience

**DOI:** 10.1101/2025.06.16.655720

**Authors:** Frederik S. Kamps, Emily M. Chen, Haoyu Du, Heather L. Kosakowski, Ariel Fuchs, Nancy Kanwisher, Rebecca Saxe

## Abstract

Sighted people rely on vision to recognize and navigate the local environment. By adulthood, human cortex contains at least three regions that respond selectively to visual scene information, but it remains unknown when or how these regions develop. One hypothesis is that scene selectivity emerges gradually in regions that initially prefer certain low-level visual features (e.g., peripheral visual input, high spatial frequencies, rectilinearity), and then exposure to the visual statistics of natural scenes drives the emergence of scene selective responses. However, both aspects of this hypothesis remain to be tested: how early scene selectivity first arises in human development, and whether it is driven by passive exposure to visual statistics. We therefore collected functional magnetic resonance imaging data from awake 2–9-month-old infants while they watched videos of real-world scenes with ego-motion, as well as faces, objects, and scrambled videos. We found stronger responses to scenes than control conditions in the location of all three scene regions. Scene-selective responses could not be explained by low-level visual properties of the stimuli, and were found in infants as young as 2-5 months old, with no evidence of age-related change. We also measured infants’ experience independently navigating (e.g., crawling), which was not necessary for the development of scene-selectivity. In sum, cortical regions are scene-selective in human infants prior to independent navigation, and after only limited exposure to visual scene statistics.

**Significance statement:** Despite extensive work on the functional organization of scene processing in the human adult visual cortex, little is known about the developmental origins of category selectivity for visual scenes. Here we used fMRI in awake human infants to discover that all three regions of the known visual scene processing system are present within the first few months of life – the youngest age yet detected. Scene-selective cortex therefore develops after only a few months of limited visual exposure, and prior to active experience using visual scene information to plan and guide independent navigation (e.g., by crawling). These findings provide a fundamental constraint on theories of how scene selectivity develops in high-level visual cortex.

## Introduction

Sighted adults adeptly use vision to recognize and find their way through the local environment, or “scene.” The foundations of this ability develop early in infancy and toddlerhood. For example, 18-month-olds use the geometry of the visual environment (e.g., the layout of walls) to navigate to a previously learned location (Hermer and Spelke, 1996), while 8-month-olds visually distinguish safe versus unsafe navigational affordances (e.g., steep versus shallow visual cliffs) (Gibson and Walk, 1960). By adulthood, the human brain contains at least three regions that respond selectively to visual scenes, and which play a causal role in scene processing and navigation (Epstein and Baker, 2019; Dilks et al., 2022): the occipital (OPA) (Dilks et al., 2013), parahippocampal (PPA) (Epstein and Kanwisher, 1998), and medial place areas (MPA; also known as the retrosplenial complex) (Maguire, 2001; Silson et al., 2016).

How does cortical scene selectivity develop? One framework (e.g., Levy et al., 2001; Arcaro and Livingstone, 2017) suggests that responses in high-level visual areas initially reflect a “protomap” of low-level visual features inherited from earlier visual cortex, with scene regions emerging in areas initially biased toward peripheral visual input, high spatial frequency content, and rectilinearity. Scene selectivity then emerges gradually through the consistent coactivation of these low-level features via passive exposure to natural scenes. We will refer to this as the “passive exposure” framework. By contrast, a second framework (e.g., Powell et al., 2018; Raz and Saxe, 2020) proposes that infants are born with systematic patterns of connectivity across the whole brain (Cabral et al., 2022; Kamps et al., 2020; Li et al., 2022), which predispose high-level visual areas to respond to functionally relevant categories early in infancy, over and above low-level visual features. Later development depends not only on passive visual exposure, but also active experience, such as with independent navigation (e.g., learning to locomote) (e.g., Held and Hein, 1963). We will refer to this as the “active experience” framework. While not mutually exclusive, these frameworks make alternative predictions about when and how cortical scene selectivity develops: i) the passive exposure framework predicts scene selectivity will emerge later, following years of exposure to visual scenes, whereas the active experience framework predicts scene selectivity will be detectable from very early on; and ii) if there is developmental change, this change will be driven by accumulating passive exposure versus active experience using scene information for navigation.

Testing these frameworks requires investigating scene selectivity in infants before, during and after their initial experiences of visual scenes and active navigation. Although numerous studies have demonstrated scene selectivity in children 5 years and older (Golarai et al., 2007; Scherf et al., 2007; Chai et al., 2010; Scherf et al., 2011; Meisner et al., 2019; Kamps et al., 2020; see also Kamps, Richardson et al., 2022), there are far fewer measurements of these cortical regions in younger children and infants. To address that gap, our lab pioneered methods using fMRI and fNIRS to study scene regions in awake infants, and found that the basic preference for scenes (relative to faces) is already established by at least 6 months (Deen et al., 2017; Powell et al., 2018), with at least one region, the PPA, showing a selective response to scenes relative to a number of control conditions (i.e., objects, faces, and bodies) (Kosakowski et al., 2022). However, it is still unclear whether responses throughout the infant scene network reflect scene selectivity versus low-level visual feature biases, how early these functions develop, and what role passive exposure and active experience may have in shaping the emergence of cortical scene selectivity.

Here we set out to address three fundamental questions: i) does scene selectivity develop throughout the cortical scene network within the first year of life? ii) do initial scene responses reflect sensitivity to low-level features, with higher-level scene selectivity emerging gradually with age, as predicted by the passive exposure framework? And iii) does the development of scene selectivity require active navigation experience? To address these questions, we scanned awake 2-9 month-old infants while viewing scenes, objects, faces, and block-scrambled versions of the scene stimuli (Figure 2b-c). To ensure quality fMRI data collection, we used methods similar to Kosakowski et al. (2022), including a custom infant head coil for improving temporal signal to noise ratio (tSNR) and strict head motion thresholds for data inclusion. To provide the most powerful test of scene-selectivity we could devise, we first conducted a pilot fMRI experiment in adults, in which we measured responses to 172 candidate scene video stimuli, and selected only the top 60 stimuli eliciting the strongest responses in adult scene regions for the current experiment (Chen, Kamps, and Saxe, 2024). Our preregistered predictions were tested using this first dataset (N=21 sessions from 17 infants; the “preregistered sample”). Then, to maximize statistical power, we combined these data with a prior sample collected using similar stimulus categories and the same head coil, by Kosakowski et al. (2022); we also report exploratory analyses from the combined datasets (N=36 sessions from 32 infants; the “full sample”).

Our preregistered predictions were as follows. First, if scene selectivity is present in infancy, then we will observe stronger responses to scenes than control stimuli in regions anatomically similar to those in adults. Second, if scene selectivity depends on passive exposure to visual scene statistics, then early emerging scene responses will be driven by low-level visual features, and scene selectivity will gradually increase with age (a proxy for passive exposure to visual scene statistics). Third, and finally, if the emergence of scene selectivity requires active experience navigating scenes, then scene selectivity will only arise in infants with locomotor experience (i.e., those who have begun crawling or walking). Our samples of usable infant data included mostly pre-locomotor infants, allowing us to test the alternative possibility that scene selectivity is present even in infants who cannot crawl or walk. Minor deviations from preregistered analysis plans are noted throughout the text.

## Results

### Estimating scene experience in infancy

Before we could test how passive and active visual experience are related to the emergence of scene selectivity, we needed to quantify infants’ visual experience with scenes over the first 9 months of life, before and after the onset of independent locomotion (crawling and walking). Surprisingly little is known about infant’s visual experience with forward-facing ego-motion through the environment, a key source of visual experience with scenes (e.g., Raudies et al., 2012; Raudies and Gilmore, 2014; Kretch et al., 2014; Gilmore et al., 2015; Petroff et al., 2025). To quantify infants’ early scene experiences, we conducted an ecological momentary assessment (EMA) study in which N = 222 parents or caregivers of infants up to age 9 months old (mean = 5.7 months) responded to 15 texts across 5 days, sent at random times between 6am and 6pm. With each text, parents reported whether their child (i) was awake, (ii) had been in motion within the last five minutes, and if so, whether the motion was (iii) self- or other-powered, (iv) forward- or backward-facing, and (v) relatively brief (i.e,. 5 minutes or less) or extended in duration (i.e., 10 minutes or more). On the first day of the study, we also asked about infants’ current locomotor status and related motor abilities.

Results are shown in Figure 1. Overall ego-motion increased steadily with age, but abundant experience was nevertheless reported prior to locomotor onset (Figure 1A). Duration of ego-motion bouts followed a similar progression (Figure 1B). Forward facing motion was present and increasing prior to the transition to self-powered motion (Figure 1C). As expected, ego-motion was initially other-powered and abruptly shifted toward self-powered motion with crawling onset (Figure 1D). Infants already faced forward from the first few months, with forward facing motion increasing sharply with age. The clear increase in reported self-motion across crawling age demonstrates that parents generally understood the questions and responded reliably, although the limited number of “yes” self motion responses under 4 months suggests at least some degree of misunderstanding or error (e.g., there is no feasible way a one month old would be capable of self-powered locomotion, per our definition). Similar results were found when accounting for children’s wakefulness at the time of response (ruling out the possibility that reported forward facing ego-motion experience early in infancy might primarily occur while infants sleep). Taken together, these data provide empirical support for several assumptions underlying our predictions: i) experience with forward-facing ego-motion begins early, prior to crawling, and accumulates steadily with age, suggesting age is a reasonable proxy for exposure to the relevant visual statistics across the first months of life; ii) human infants have little-to-no experience with active navigation (via self-powered motion) prior to five months of age; and iii) forward-facing ego-motion through scenes occurs before and independently of active locomotion, so effects of active locomotor experience are potentially separable from those of passive visual exposure to optic flow patterns characteristic of scenes.

**Figure 1.**
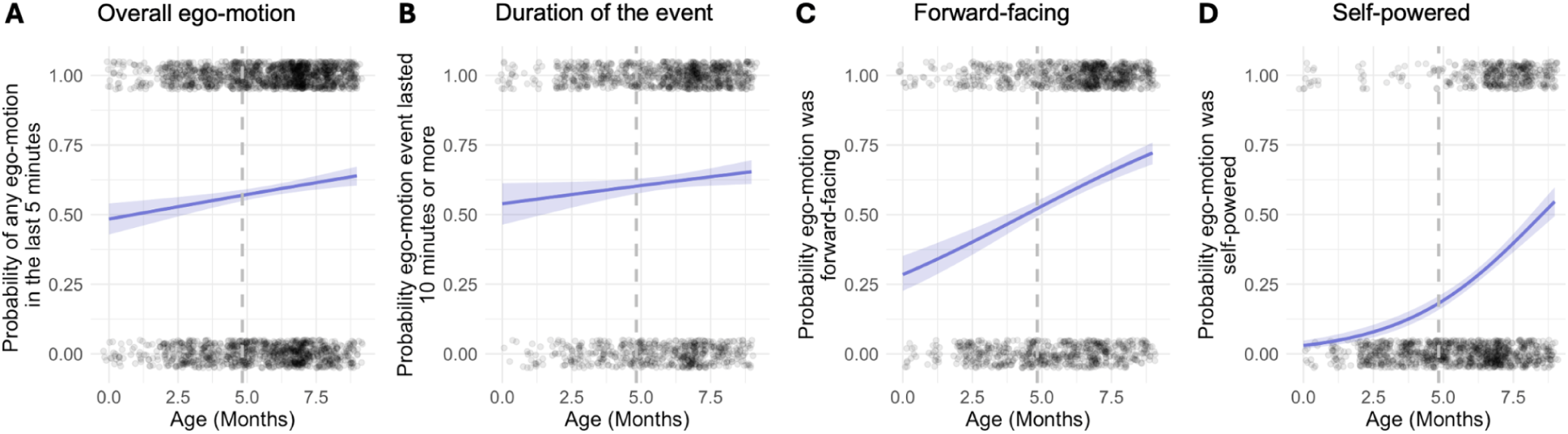
*EMA study of experience with ego-motion through the environment across the first 9 months.* (A) Proportion of survey responses indicating that yes their child had experienced any kind of ego-motion in the past 5 minutes, as a function of age. (B) Proportion of the “yes” ego-motion responses presented in (A) where the parent also indicated the child had been in motion for a period of 10 minutes or more, as a function of age. (C) Proportion of “yes” ego-motion responses where parents indicated the child was facing forward during the ego-motion, as a function of age. (D) Proportion of “yes” ego-motion responses where parents indicated the motion was self-powered (rather than other-powered), as a function of age. For all plots, blue lines indicate the fit of a logistic regression model with shaded confidence intervals. Dots jittered around 0 and 1 indicate individual survey responses. Gray vertical lines indicate the youngest age at which any parent indicated their child was able to crawl.

How much experience do infants gain in the first two to nine months of life? We also used the EMA data to estimate infants’ total (lifetime) hours of ego-motion experience (details of calculations are provided in the supplemental materials). We estimate that infants ages 2 to 9 months old (the age range of our infant fMRI sample) have experienced approximately 146-1427 hours of awake ego-motion, and 62-1223 hours of forward-facing ego-motion (i.e., as depicted in our scene stimuli). A 4.9 month old (the median age of the full sample) is estimated to have experienced 522 hours of ego-motion, and 338 hours of forward facing ego-motion. Notably, these estimates are likely an upper bound for visual scene experience, given that many relevant aspects of basic visual perception are limited in the first few months, including visual acuity (impacting high spatial frequency and rectilinearity experience) and the extent of peripheral vision (Daw and Daw, 2006; Atkinson, 2002).

### The cortical scene network is scene selective in infancy

Having established that effects of passive exposure and active experience are potentially dissociable, we next tested whether scene responses can be detected using fMRI in awake, 2-9 month old infants. To do so, we conducted a standard subject-specific functional region of interest (ssfROI) analysis. ssfROIs were defined within previously described anatomical parcels, or “search spaces”, that capture the typical spatial locations of the OPA, PPA, and MPA in adults (Julian et al., 2012) (Figure 2A). Given the imperfect registration of adult anatomical templates to infant functional data, each parcel was slightly increased in size, following the approach used in Kosakowski et al. (2022). For each potential infant subject, we first defined “subruns” as portions of the originally acquired data during which infants were awake and still (see Methods for full details on subrun definition). The preregistered dataset included an average of 524.9 volumes (26.3 minutes) per infant (range: 199-1542), and the Kosakowski et al. (2022) dataset included an average of 533.9 volumes (26.7 minutes; range: 207-1182 volumes) per infant. To estimate scene selectivity for each session, we used a leave-one-subrun-out procedure to define each ssfROI and independently test its response profile. ssfROIs for scene regions were defined as the top 5% of voxels in each parcel responding to scenes > control stimuli (the average of faces, objects, and scrambled for the preregistered sample, or faces, objects, and bodies for the full sample) stimuli. Responses to the experimental conditions (Figure 2B-C) in the ssfROI were measured using the left out subrun, and results were averaged across all possible splits of subruns.

**Figure 2.**
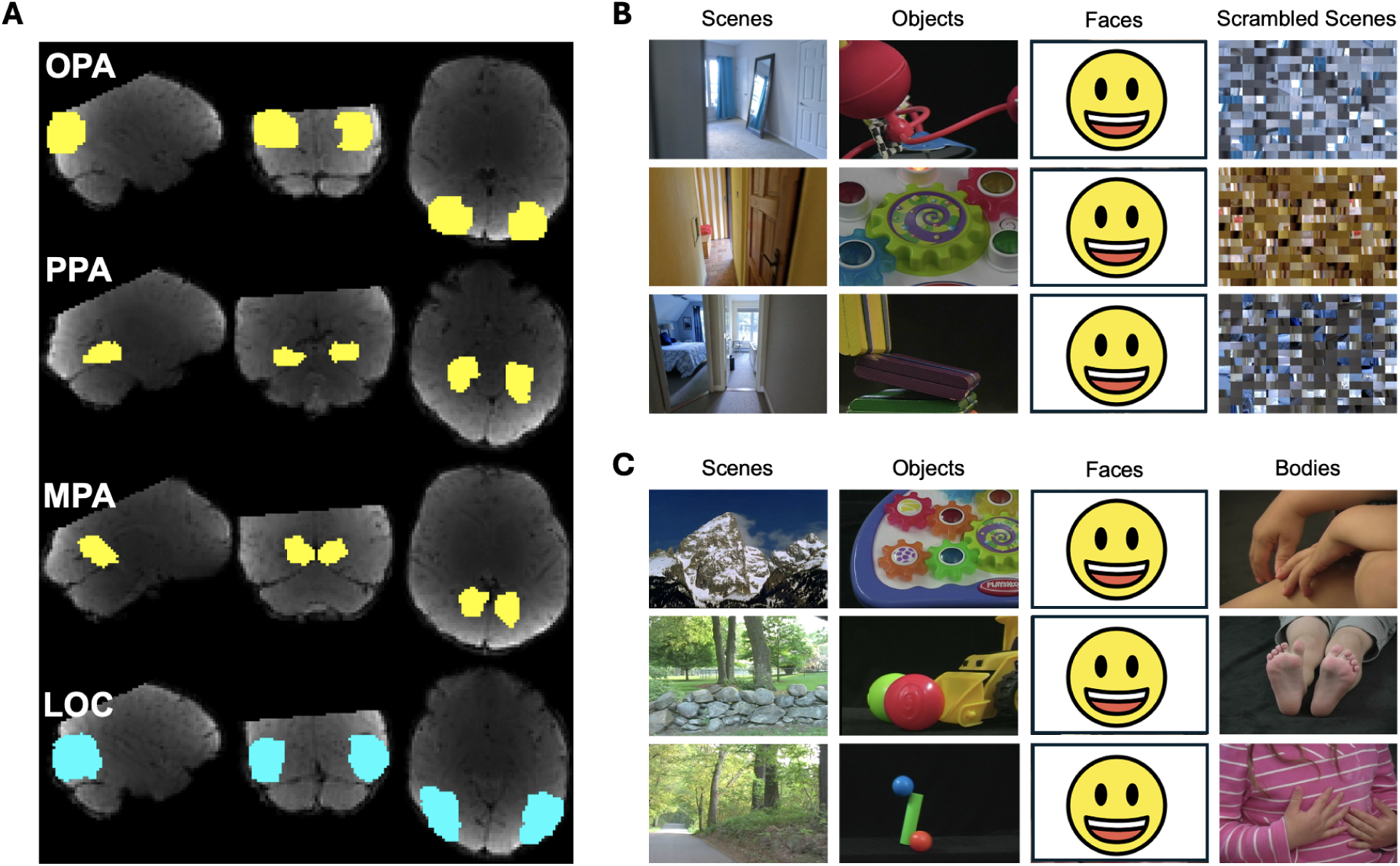
*ssfROI search spaces and experimental stimuli.* (A) Search spaces for each region, shown in infant template space. Subject-specific functional ROIs were identified as the top 5% of voxels within each search space, based on the contrast of [scenes] - [objects, faces, scrambled] (for scene regions in yellow) or [objects, faces] - [scenes, scrambled] (for object regions in blue). Note that although some parcels included off-brain voxels, due to the increased size of the parcels, and the imperfect registration of adult parcels to infant template space, functional data were carefully masked to ensure that no off-brain voxels were selected via the ssfROI procedure. (B) Example frames from the dynamic movie stimuli for each of the four conditions in the preregistered dataset. (C) Example frames from the dynamic movie stimuli for each of the four conditions in the dataset from Kosakowski et al. (2022). Face images redacted per BioRxiv policy.

Results of the ssfROI analysis are shown in Figure 3. We began by comparing responses to scenes versus the average of all control stimuli in each infant scene region. For the preregistered sample, we found greater responses to scenes than control stimuli in OPA (β = 8.42, t_(61.49)_ = 2.62, p = 0.006), PPA (β = 2.28, t_(64.94)_ = 1.91, p = 0.03) and MPA (β = 4.19, t_(59.80)_ = 2.58, p = 0.006), with no significant region x condition interaction for any pair of scene regions (all F’s < 1.30, all p’s > 0.27). Similar results were found for the three scene regions in exploratory analyses of the full sample (all β’s > 3.11, all t’s_(104.66)_ > 2.72, all p’s < 0.004; no significant region x condition interaction for any pair of scene regions: all F’s < 1.60, all p’s > 0.19). To provide a stricter test of scene selectivity, we also compared responses to scenes with each individual control condition (noting that we had less power to detect these effects, given that each control condition was shown 1/3 less often than the scene condition in the preregistered dataset). For the preregistered dataset, an exploratory omnibus analysis considering data from all three scene regions (with a factor for region) revealed greater responses to scenes than each of the three control conditions (all β’s < -8.13, all t’s_(224.01)_ < -1.91, all p’s < 0.03). Similarly, for the full dataset, this analysis (now including an additional factor for experiment) revealed greater responses to scenes than to each of the four control conditions (all β’s < -7.06, all t’s_(390.91)_ < -2.10, all p’s < 0.02). Results for these comparisons tested separately in each scene region are summarized in Figure 3 (B and D) and detailed in the supplemental materials. Taken together, these results suggest that scene selectivity is present throughout the cortical scene network within the first year of life.

**Figure 3.**
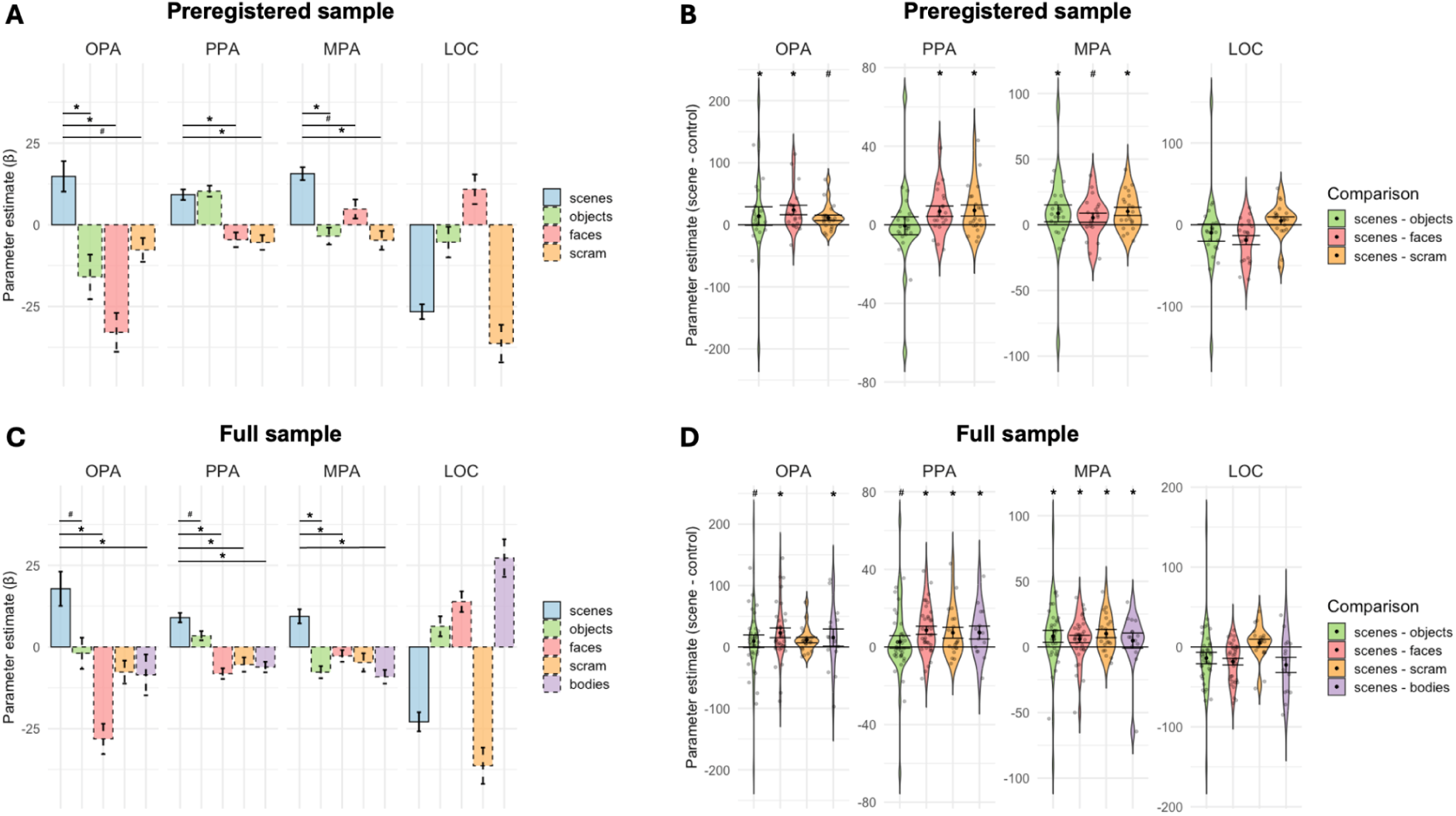
Scene-selective responses are detectable in 2-9 month old infants, and functionally dissociable from responses in nearby object-preferring cortex. (A) Data from the preregistered sample of infants only. Bar plots depict the mean response to scenes versus each individual control condition (objects, faces, and scrambled scenes). Dashed lines on control conditions reflect the fact that these conditions were sampled 33.3% as often as the scene condition (in the preregistered dataset only). Error bars indicate standard error of the mean. (B) Violin plots depicting the difference in response to scenes minus each control condition. Error bars indicate standard error of the mean. Gray points indicate individual subjects. (C and D) Same as A and B, respectively, but now plotting data from the full sample of infants. Note that the full sample of infants included infants from the preregistered dataset, who viewed scenes, objects, faces, and scrambled scenes, as well as infants from scenes, objects, faces, and bodies; accordingly, fewer infants contributed to the estimates of scrambled scenes and bodies. Asterisks indicate significant results from the linear mixed effect model (* = p < 0.05, # = p < 0.10).

An exploratory group-level whole-brain analysis confirmed that the spatial organization of scene responses is qualitatively similar to that found by adulthood. Figure 4 shows results from a group random effects analysis for the contrast of scenes vs. the average of all control conditions using the full sample (N=32 subjects). At low (uncorrected) thresholds, greater activation to scenes is found in areas consistent with the spatial location of adult scene regions, while greater activation to the control conditions is found in areas consistent with the spatial location of adult object and face regions. However, group-level activations in infant scene regions were not significant after correcting for multiple comparisons.

**Figure 4.**
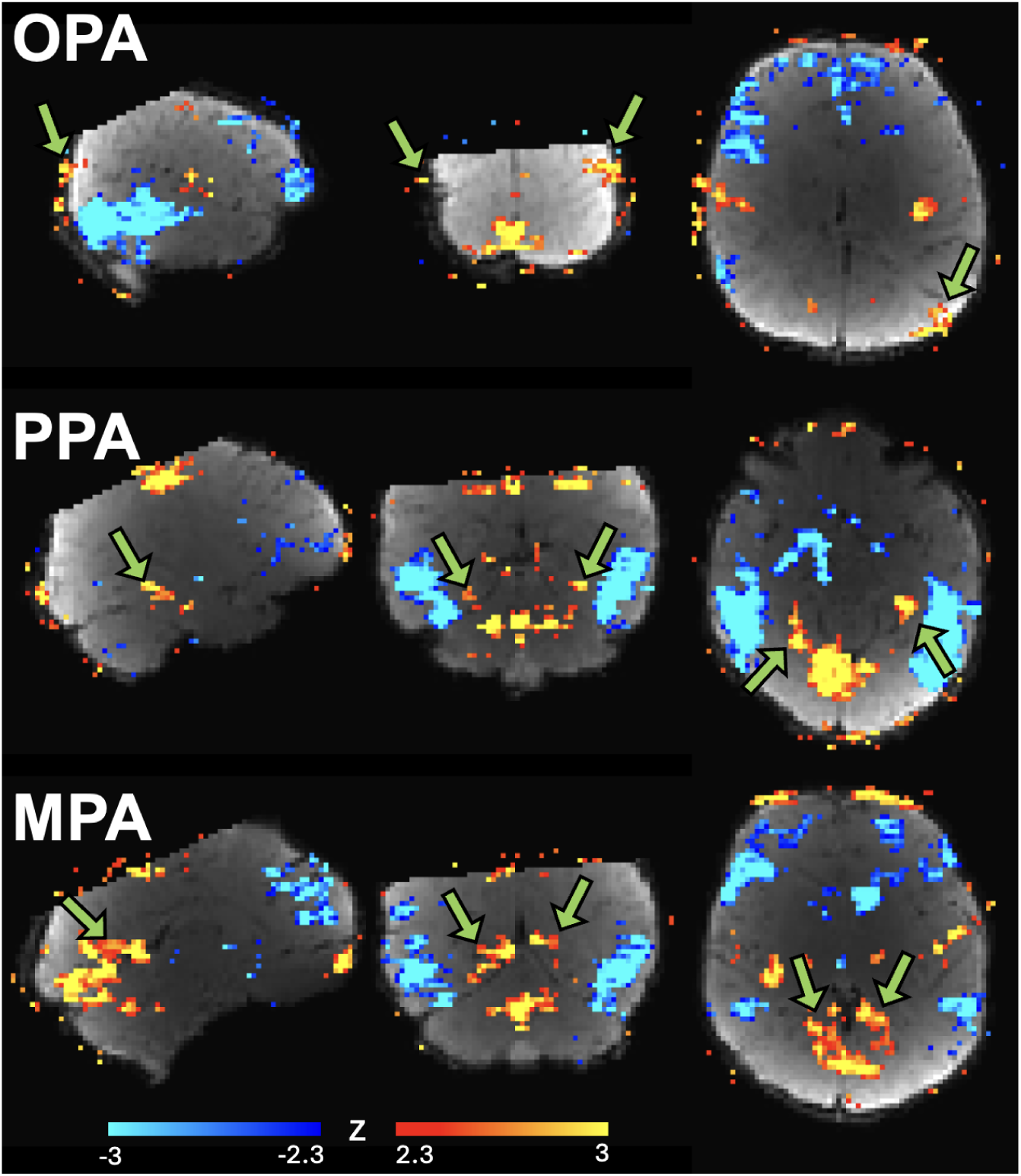
Whole-brain group random effects analysis. Maps show activation across subjects (N=32) for the contrast of scenes vs. the average of all control conditions, with regions preferring scenes in warm colors, and regions preferring the control conditions in cool colors. Data are shown at a low, uncorrected threshold (p < 0.01; transformed to z for visualization). Activation in scene regions was not significant after accounting for multiple comparisons. Results are displayed in infant template space (see methods).

### Infant scene cortex is functionally dissociable from nearby object-preferring cortex

By adulthood, scene areas are functionally distinct from nearby high-level visual areas sensitive to objects (Park et al., 2011; Kravitz et al., 2011). To test for this division of labor in the infant cortex, we compared the functional profile of the infant scene regions with that in a nearby area in lateral occipital cortex (LOC) sensitive to objects (i.e., any stimulus with a visual contour, including objects and faces, but not intact or scrambled scenes). LOC was identified using an adult parcel, with ssROIs selected based on the contrast of either [objects and faces] minus [scene and scrambled scenes] for the preregistered dataset, or objects minus scenes for the Kosakowski et al. (2022) dataset (we could not use the same contrasts across datasets due to differences in conditions tested and amount of data per condition). LOC responses across the conditions are shown in Figure 3. Beginning with the preregistered dataset, we found a significant region x condition interaction for each scene area compared individually to LOC (all β’s < 3.18, all t’s_(144.00)_ < 1.75, all p’s < 0.042). Similar results were found using the full sample (now including a factor for experiment), with each scene region again showing a significant region by condition interaction, relative to LOC (all β’s > 4.85, all t’s_(249)_ > 3.22, all p’s < 0.001). Scene-selective cortex is therefore functionally dissociable from nearby object-preferring regions within the first nine months of life. This dissociation of responses across regions also suggests that the pattern of responses in infant scene cortex cannot be explained by general differences in attention, arousal, or basic visual properties (e.g., eyes open vs. closed, luminance) across the experimental conditions.

### Infant scene responses are not explained by low-level visual features

A key prediction of the passive exposure framework is that early-emerging responses in scene regions reflect biases for peripheral visual input and low-level visual features (including high spatial frequency content and rectilinearity), rather than a higher-level, category-selective response. To test this prediction, we quantified the mean high spatial frequency, rectilinearity, and (peripheral) motion energy for each of our stimulus categories using standard approaches (see Methods and Supplemental Figure 1). Given that infants may have looked anywhere in our stimuli, we measured motion energy both in the entire video, and also specifically in the periphery of the video. We then fit exploratory linear mixed effect models with two factors: a categorical model of scene selectivity (i.e., with scenes coded as 1 and all other conditions as -1) and a low-level feature model (i.e., the normalized mean values of each feature across conditions, tested individually against the category model). To maximize power, we included data from all scene regions (results from individual regions are reported in the supplemental materials). For the preregistered dataset, we found a significant effect of category tested against all four low-level feature models (all β’s > 4.38, all t’s_(225.00)_ > 2.34, all p’s < 0.02), with none of the four low-level features explaining significant variance after accounting for the category model (all β’s < 1.31, all t’s_(225.00)_ < 0.80, all p’s > 0.42). The same pattern of results was found in the full dataset, with a significant effect of category (all β’s > 4.53, all t’s_(392.63)_ > 2.89, all p’s < 0.005), but no significant effect of any low-level feature (all β’s < 1.28, all t’s_(392.63)_ < 0.96, all p’s > 0.37).

These results suggest that the functional profile of infant scene regions is better explained by scene selectivity than any single low-level feature. Do low-level features play any role in driving infant scene regions responses, beyond this scene selective profile? To explore this question, we tested the low-level feature models on the non-scene control conditions only, following similar comparisons of non-scene stimuli in adult scene regions (Nasr et al., 2014; Bryan, Julian et al., 2016). However, none of the four low-level features predicted significant variance in infant scene region responses to the control categories, in either the preregistered dataset (all β’s < 1.40, all t’s_(165.03)_ < 0.77, all p’s > 0.44) or the full dataset (all β’s < 1.36, all t’s_(287.33)_ < 0.96, all p’s > 0.33). We also considered the possibility that a combination of low-level visual features might be required to drive responses in infant scene regions, rather than any single feature. However, a model based on a weighted sum of the low-level features (see Supplemental Materials for details on how weights were derived) also failed to explain significant variance in infant scene regions, relative to a categorical model. Taken together, we find no evidence for the hypothesis that infant scene responses reflect biases for low-level visual features.

### No evidence of age-related change

A second prediction from the passive exposure framework is that scene-selective responses develop gradually with age, through accumulating exposure to visual scene statistics across months if not years. Testing this prediction here across ages 2 to 9 months old, we found no interaction of age and scene selectivity (i.e., scenes vs. avg(control conditions)) in any of the three regions (preregistered sample: all β’s < 1.05, all t’s_(63.94)_ < 0.87, all p’s > 0.39; full sample: all β’s < 2.25, all t’s_(103.49)_ < 1.87, all p’s > 0.06). We also failed to detect an age x scene selectivity interaction in an exploratory omnibus regression model including data from all three scene regions (preregistered sample: β = 0.28, t_(227.28)_ = 0.16, p = 0.87; full sample (β = 0.92, t_(392.64)_ = 0.69, p = 0.49). To test whether scene-selectivity is present even in the youngest infants, we median split the full sample of sessions based on age at acquisition, yielding a younger sample of 19 sessions from 18 infants; mean age = 3.71 months, range = 2.07-4.93 months). An exploratory omnibus analysis (across regions, with a factor for region) found greater responses to scenes than control stimuli (β = 3.38, t_(206.68)_ = 1.80, p = 0.04) (results from the same analysis in the preregistered sample, as well as from individual regions, are reported in supplemental materials). Taken together, these analyses failed to reveal evidence of age-related change in cortical scene responses during the first nine months, and rather suggest that scene selective responses are relatively stable across the first nine months, emerging as early as 2-5 months old.

### Scene responses develop prior to onset of independent navigation

Is active experience using scene information for self-guided locomotion required for the development of scene selectivity? If the development of scene selectivity does not depend on active navigation experience, then scene regions will be present even in pre-locomotor infants. To test this prediction, we analyzed the sample of N=16 sessions from 13 infants from the preregistered dataset who had never independently locomoted at the time of acquisition (by crawling, walking, or any other means of moving independently to a location more than 3 feet away, per parent report at the time of scan) (mean age at time of session = 4.70 months, range = 2.07-7.89 months). We compared responses to scenes versus the average of all control stimuli in each infant scene region (Figure 6). This analysis revealed greater responses to scenes than the control conditions in OPA (β = 9.40, t_(45.97)_ = 2.80, p = 0.004) and MPA (β = 3.99, t_(45.72)_ = 2.33, p = 0.01), although this comparison did not reach significance in PPA (β = 1.93, t_(49.43)_ = 1.48, p = 0.07). Given the limited sample size of pre-crawling infants in the preregistered dataset, we also analyzed the full sample of infants. Locomotor status was not assessed directly for infants in Kosokowski et al. (2022). Accordingly, for these infants, crawling status was estimated using our EMA data. We identified the youngest age at which any infant in the EMA sample (N= 222) was reported to be able to locomote: age 4.83 months. There were 9 infants younger than 4.83 months old in the Kosokowski et al. (2022) dataset, yielding a total sample of 25 sessions from 22 pre-locomotor infants (mean age at time of session = 4.46 months, range = 2.07-7.89 months). Analysis of this larger pre-locomotor sample revealed greater responses to scenes than control conditions in OPA (β = 7.28, t_(72.90)_ = 2.25, p = 0.01), PPA (β = 2.78, t_(76.48)_ = 2.54, p = 0.007) and MPA (β = 2.67, t_(71.22)_ = 1.85, p = 0.03). Taken together, these results show that scene-selectivity emerges throughout the scene network prior to locomotor experience.

**Figure 5.**
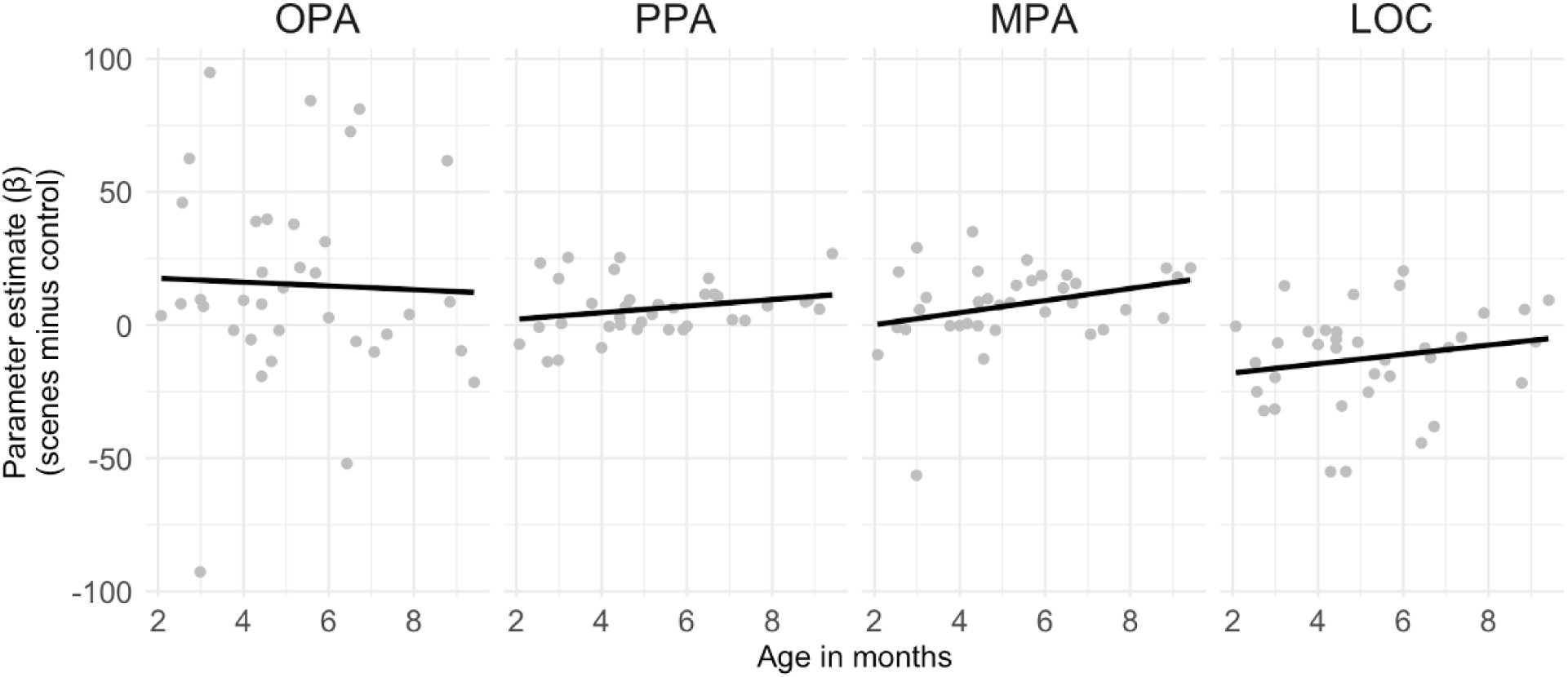
*No evidence of age-related change in scene-selectivity*. Each plot depicts the difference in response to scenes minus the average of the control conditions as a function of age, using the full sample of infants. Gray dots indicate individual infants. No region showed a significant effect of age.

**Figure 6.**
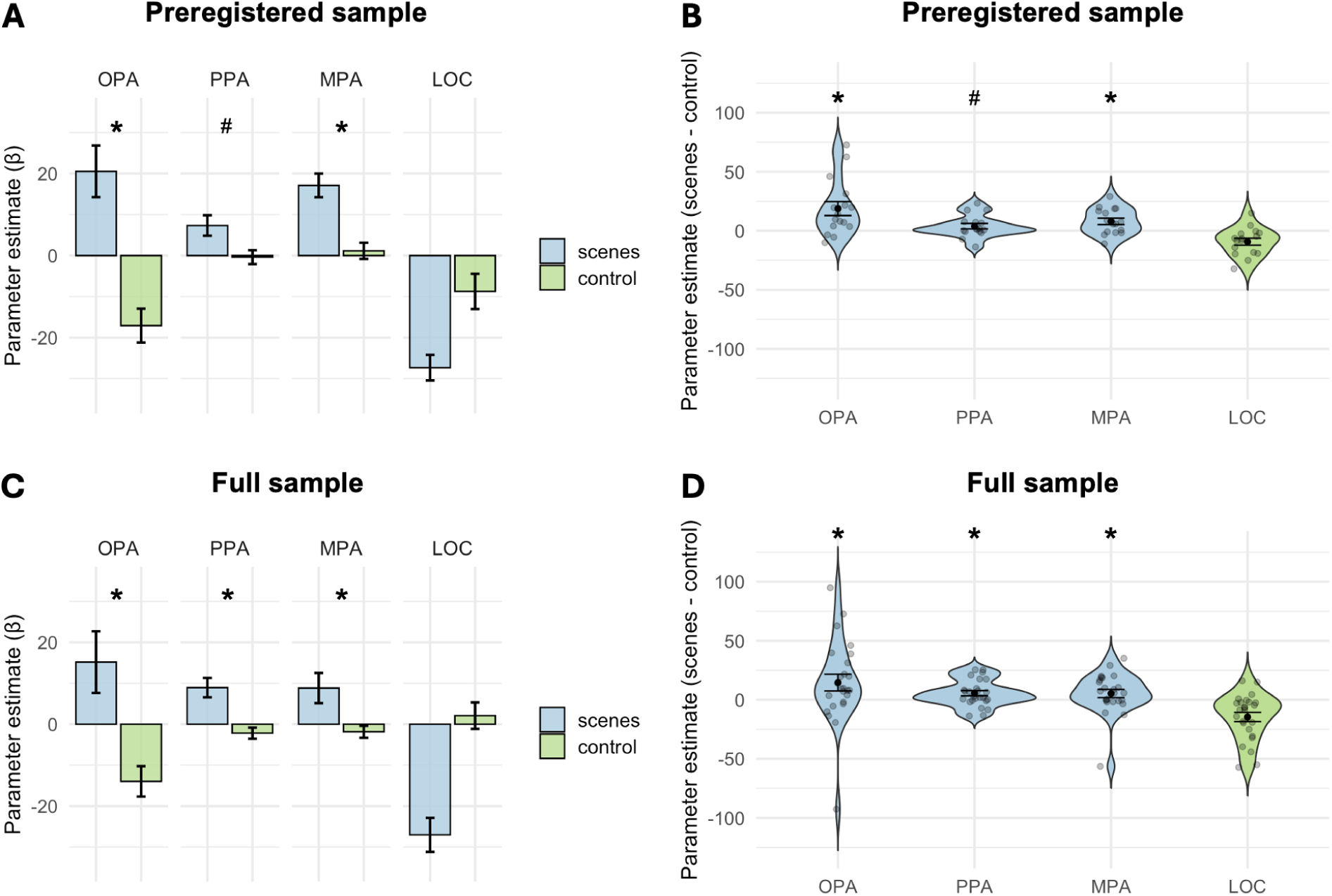
*Scene responses are present in pre-locomotor infants.* (A) Data from the preregistered sample of pre-locomotor infants only. Bar plots depict the mean response to scenes versus the average of the control conditions (objects, faces, and scrambled scenes). Error bars indicate standard error of the mean. (B) Violin plots depicting the difference in response to scenes minus the mean of the control conditions. Error bars indicate the standard error of the mean. Gray dots indicate individual subjects. (C and D) Same as A and B, respectively, but now plotting data from the full sample of pre-locomotor infants.

## Discussion

Here we found that scene selectivity develops in all three known regions of the cortical scene processing network, in similar locations to adults, by just 2-9 months of age. Responses in infant scene regions were i) dissociable from those in nearby object-preferring cortex, ii) not easily explained by confounds in low-level visual properties (e.g., peripheral visual stimulation, high spatial frequency content, rectilinearity), and iii) found even in the subsample of pre-crawling infants, as well as the subsample of infants less than five months old. Selective responses to scenes therefore do not require years of experience to develop, but rather emerge rapidly after only a few months of visual exposure, and prior to active experience using scene information to guide independent navigation.

These findings build on prior awake infant fMRI studies showing preferential responses scenes versus other categories, particularly in the PPA (Deen et al., 2017, Kosakowski et al., 2022), by revealing that selective responses are in fact present across all three regions of the cortical scene network in infancy. We speculate that the success of the current study in finding scene selective responses in all three regions may be attributed to the increased size of the sample of data collected using the newer custom head coil (Ghotra et al., 2021) and higher-resolution EPI sequences, relative to the datasets used in our previous studies (Kosakowski et al., 2022; Deen et al., 2017). Another possibility is that aspects of the current design, such as using dynamic stimuli that maximally activate scene regions in adults (Chen, Kamps, and Saxe, 2024), might have contributed to our success–although we found no reliable difference in scene selectivity between the preregistered and Kosakowski et al. (2022) datasets. Combining the two samples increased the statistical power of our analyses, and revealed significantly higher responses to scenes than each of four control conditions tested (objects, faces, scrambled scenes, bodies) in all three regions. In the smaller preregistered sample alone, a few of these comparisons were non-significant trends, confirming the value of a larger sample.

The presence of scene-selectivity in 2-9-month-old infants places new constraints on the passive exposure framework for the development of category-selective visual areas (Acaro and Livingstone, 2021). Indeed, here we find scene selectivity is already present earlier than the current earliest evidence for retinotopy in awake human infants (Ellis et al., 2021). Accordingly, if passive exposure drives the initial development of scene responses, this process must occur within the first hundred days of visual experience. Recent computational modeling suggests that such a rapid timeline is plausible: for example, DNNs trained with just 200 hours of head-camera footage from a single child achieve 70% of the accuracy of a high-performance ImageNet-trained model (Orhan and Lake, 2023). However, this work sampled only older infants’ visual diets, and did not account for the limited nature of very young infants’ vision. In the first five months, many relevant visual processes are still immature, including visual acuity (and thus sensitivity to high-level spatial frequencies and rectilinearity), peripheral visual input, depth and distance perception, and oculomotor control (Atkinson, 2008; Daw and Daw, 2006). Accordingly, our results are also consistent with the hypothesis that very little visual exposure is necessary to support the initial emergence of scene selectivity, consistent with evidence of scene responses in the congenitally blind (Wolbers et al., 2011; He et al., 2013). Subsequently, visual exposure may support a gradual process of refinement of scene representations across years of childhood visual experience (e.g., Gomez et al., 2019), although evidence for age-related change in scene regions across childhood is inconsistent (e.g., see Golarai et al., 2007; Scherf et al., 2007; Scherf et al., 2011; Jiang et al., 2014; Meisner et al., 2019; Kamps et al., 2020; Kamps, Richardson, et al., 2022; Jung et al., 2024). The finding that scene regions are functionally dissociable from nearby object regions early in infancy is consistent with evidence that connectivity underlying high-level visual areas may be present from birth, potentially guiding the development of category selectivity from early on (Arcaro et al., 2017; Kamps et al., 2020; Li et al., 2020; Cabral et al., 2022).

Our findings do rule out the hypothesis that the development of scene selectivity requires active functional experience using visual scene information to guide locomotion through the environment. Scene selectivity was observed in infants with no locomotor experience. This finding is a key result of the current study, consistent with but more definitive than prior observations of scene selective responses in the infant cortex (Deen et al., 2017; Powell et al., 2018; Kosakowski et al., 2022). Active locomotor experience may still refine visual scene representations later in childhood. Growing evidence suggests OPA, PPA, and MPA ultimately extract different representations and serve distinct functional roles within scene processing (Dilks et al., 2023). Future research should test whether these divisions of labor are already present in infancy, or mature more slowly (e.g., Kamps et al., 2020; Kamps, Rennert et al., 2023; Jung et al., 2024), and whether their development requires specific kinds of visual experience.

In conclusion, here we show that scene selectivity is present throughout the cortical scene network in infancy, years earlier than previously observed, and prior to experience with independent navigation. These findings provide a fundamental datapoint for theories of how these regions develop, and more broadly highlight the potential of cortical scene processing as a test case for understanding the relative roles of active experience and passive exposure in shaping the development of the human brain.

## Methods

### Participants

Subjects were healthy human infants ages 2-9 months old. This age range was chosen because it is the earliest age at which scanning awake human infants is feasible (Kosakowski et al., 2022; Deen et al., 2017; Kosakowski et al., 2024). Participants were recruited via online ads posted on social media sites (Facebook and Instagram), local posters, and word of mouth. The final sample size depended on several thresholds for inclusion based on data quality and quantity, outlined in detail below. Infants were invited to return for multiple sessions; sessions conducted less than 31 days apart were treated as a single session and analyzed together, while sessions conducted more than 31 days apart were analyzed separately. These factors lead to final sample sizes of 21 sessions from 17 infants (mean age at time of session = 5.40 months, range = 2.07-9.11 months) for the preregistered sample, and 36 sessions from 32 infants in the full sample (mean age at time of session = 5.25 months, range = 2.07-9.41 months). In addition to the fMRI experiment, parents of infants in only the preregistered sample completed an in-lab demographic questionnaire, as well as a subset of twelve questions taken from the Peabody Developmental Motor Scales, selected to identify experience with independent navigation, from belly crawling to walking with and without support. Parents were also asked to report directly whether their infant has ever successfully crawled or walked. Specifically, we asked parents whether, and if so, how their child is capable of locomoting toward a goal – by any means, including any form of crawling – more than three feet away. For navigating infants, we further collected a calendar date of crawling or walking onset (per parent report).

### Stimuli

Stimuli for the preregistered dataset are available at https://osf.io/vk7aw/. Stimuli for the Kosakowski et al. (2022) dataset are available at https://osf.io/jnx5a/, and are described in detail in the paper. For the preregistered dataset, scene stimuli were chosen based on the results of a pilot fMRI experiment in five adult participants. In this pilot experiment, responses were measured to 172 candidate scene movies using an event-related design (Chen, Kamps, and Saxe, 2024). Candidate scene movies included 9 representative scene movies taken from two previous infant fMRI studies (Deen et al., 2017; Kosakowski et al., 2022; Kosakowski et al., 2024), as well as 163 new scene movies, which we recorded in varying locations around the world. The set of candidate scene stimuli included considerable variation in the nature of ego-motion through the space, the spatial layout of the scene (e.g., open vs. closed), the materials depicted (e.g., manmade/indoor vs. natural/outdoor), and the affordances of the space (e.g., ability to navigate to the left vs. right). As control stimuli, we also included 3 face movies, 6 “baseline” movies, and 9 object movies, all taken from a previous infant fMRI experiment (Kosakowski et al., 2022; Kosakowski et al., 2024). In all three adult scene regions, we observed stronger responses to the new scene movies compared to control stimuli, as well as considerable variability in the response to the different scene movies, with the previously used scene movies exhibiting weaker responses relative to the new scene movies. Indeed, many of the scene videos used in the previous infant fMRI experiments elicited responses that were similar to or only slightly stronger than the responses to objects, whereas many of the new candidate scene videos elicited responses that were 1.7-2.5 greater than the older scene movies (depending on the region).

For the preregistered experiment, we then selected the top 60 scene videos from this analysis based on responses in the OPA to maximize the chance of identifying scene-selective responses in infants (notably, univariate responses to the candidate videos were highly correlated across OPA, PPA, and MPA, such that top videos for OPA also strongly activated the adult PPA and MPA) (Chen, Kamps, and Saxe, 2024). In addition to scene videos, infants also viewed stimuli from three control categories to provide a test of scene-selectivity, including objects, faces, and scrambled scenes. Face videos were taken from Pitcher et al. (2012) and depicted only the faces of infants and children against a black background. Object videos were taken from Pitcher et al. (2012) and Kamps et al. (2022) and depicted colorful, inanimate toys and household objects against a black background. Although scene information was minimized in the object videos due to the use of a black background, special care was taken to select only those object videos that did not even imply scene information. For example, an object rolling along a surface might imply the presence of a ground plane, and therefore the presence of a basic scene layout. Finally, scrambled scene videos were created out of the top 20 scene videos by dividing the scene videos into 15x15 blocks, and scrambling the position of those blocks, thus disrupting the coherent sense of scene content while preserving many local, low-level visual features present in the scene. We used 20 videos for each control condition, and 60 different videos for the scene condition, since scene blocks were shown three times as often as control blocks, thus matching visual adaptation between conditions. Finally, to further attempt to maximize BOLD responses and increase infant attention, each video was shown for 2.7s, followed by a 300ms still image from the same category (but not taken from the same videos), following the design of Kosokowski et al. (2022; 2024). Scene, face, and object still images were taken from a variety of sources (Kosakowski et al., 2022; Kamps et al., 2016; Walther et al., 2011; Park et al., 2015), and scrambled scene still images were created from the still scene images using the same block-scrambled approach described for the scrambled scene videos above.

### Low-level visual feature analysis

We analyzed three kinds of lower-level visual feature content previously showed to drive responses in adult scene regions and hypothesized to drive scene region responses early in development, including high spatial frequency content (Levy et al., 2001), rectilinearity (Nasr et al., 2014), and peripheral visual input (operationalized as motion energy, particularly in the periphery of the video stimulus) (Levy et al., 2004). Visual statistics were analyzed for each video or image stimulus (Figure 5). For video stimuli, each frame was extracted from each video. Frames and still images were converted to grayscale and normalized. To obtain spatial frequency information, we computed a Fourier transform on each image. High-spatial frequency content was extracted from each frame of each stimulus using the methods and cut-offs described by Rajimehr et al. (2011), and averaged across all images within a category. Each category value was then normalized relative to the baseline value. For rectilinear information, angled Gabor filters (90 and 180) with four different spatial frequencies (1, 2, 4, and 8) were applied to each pixel to assess the amount of angular content. Rectilinear values were averaged across pixels and blocks. To obtain motion energy information, we used the approach developed by Nishimoto et al. (2011), in which grayscale videos were passed through a bank of three-dimensional spatiotemporal Gabor wavelet filters. For each video, motion energy signals were compressed by a log-transform and summed across the bank of filters, yielding an estimate of motion energy per video. To estimate motion energy specifically in the periphery of the video, we applied a white rectangle over the center of the video (corresponding to 6.9 x 9.1 degrees visual angle), such that only the peripheral content of the video remained visible, and computed motion energy on the resultant videos.

### Design

Stimuli were presented in a block design at 13.7 x 18.1 degrees visual angle, with six 3s video + still image pairs presented per block, yielding 18s blocks. For only the preregistered dataset, the block sequence alternated between scenes and control blocks, such that scene blocks were shown every other block, and the three control blocks were shown in a pseudorandom sequence such that all three control conditions were always shown before any scene condition was repeated. This design was chosen to ensure adequate sampling of scene activation in any given subrun. A block of colorful attention-getter stimuli, with accompanying sounds, was also shown every 8 blocks. A run began as soon as the infant was in the scanner and videos played continuously for as long as the infant was engaged and awake. If infants fell asleep, we continued collecting resting state data for at least 10 additional minutes. Similar methods were used for the Kosakowski et al. (2022) dataset, except that scene blocks were presented in approximately equal proportion to control blocks, and the baseline condition consisted of curvy, abstract movies rather than attention getters with accompanying sounds. Potential differences in baseline activation between the two datasets were accounted for via a factor for experiment in all linear mixed effects models of the full dataset.

### Data Collection

Infants were swaddled if the caregiver thought it would be useful. A parent or researcher went into the scanner with the infant while a second adult researcher (“scanner buddy”) stood outside the bore of the scanner, while maintaining a view of the infant’s face through the bore. The scanner buddy monitored infant attention and indicated to a researcher in the control room when the infant was not looking at the stimuli. The researcher in the control room then recorded these TRs for exclusion during preprocessing. Infants heard lullabies (https://store.jammyjams.net/products/pop-goes-lullaby-15) played through custom infant headphones for the duration of the scan.

### Custom Head Coil and Equipment

Infants were scanned in a 32-channel head coil designed to comfortably cradle infants while wearing custom infant MR-safe headphones (Ghotra et al., 2021). Infant headphones attenuated scanner noises (attenuation statistics reported in Ghotra et al., 2021) and allowed infants to listen to music at a comfortable volume for the duration of the scan. An adjustable coil design increased infant comfort, accommodated headphones, and suited a variety of infant head sizes. The infant head coil and headphones were designed for a Siemens Prisma 3T scanner and enabled the use of a standard trajectory EPI with 43-44 near-axial slices (repetition time, TR = 3s, echo time, TE = 30ms, flip angle = 90, field of view, FOV = 160 mm, matrix = 80x80, slice thickness = 2 mm, slice gap = 0.0-0.4 mm).

### Data Selection (subrun creation)

To be included in the analysis, data were required to meet the criteria for low head motion that was reported by Deen et al. (2017) and Kosakowski et al. (2022). Data were cleaved between consecutive time points having more than 2 radians or 2 millimeters of motion, creating “subruns” which contain at least 24 consecutive low-motion volumes. All volumes included in a subrun were extracted from the original “run” data (a “run” began when the subject went into the scanner and ended when the subject fell asleep or became fussy) and combined to create a new NIfTI file for each subrun. Event files were similarly updated for each subrun. Volumes with greater than 0.5 radians or millimeters of head motion between volumes were scrubbed. To be included in the ssfROI analysis, subjects were required to have at least two usable subruns of length 96 volumes each (one to choose voxels based on the relevant contrast, and the other to independently extract response magnitudes from the selected voxels). We originally preregistered a lower threshold for data inclusion, requiring just two usable subruns of 24 volumes each. However, upon analyzing the data from Kosakowski et al. (2022) in a pilot analysis focused on responses in the infant fusiform face area, we discovered that inclusion of infants with such low data quantity led to noisier selectivity estimates of face selectivity. We therefore chose to require 96 volumes in each split-half of the data, following the approach used in Kosakowski et al., 2022. For subjects with only one subrun, the originally selected subrun was split in half (discarding the middle two volumes), and each half was reevaluated for meeting the subrun criteria described above. If both halves were usable, the subrun was split; if not, the subject was discarded. After carrying out these procedures, infants in the preregistered dataset had an average of 524.9 volumes (i.e., 26.3 minutes) per infant (range: 199-1542), with an average of 71.2% (range: 39.3-92.7%) of volumes below the motion threshold of 0.5mm or degrees). Infants in the Kosakowski et al. (2022) dataset had on average of 533.9 volumes (i.e., 26.7 minutes; range: 207-1182 volumes) per infant, with an average of 61.5% (range: 32.8-86.1%) of volumes below the motion threshold of 0.5 millimeters or degrees).

### Preprocessing

Each subrun was processed individually. First, an individual functional image was extracted from the middle of the subrun to be used for registering the subruns to one another for further analysis. Then, each subrun was motion corrected using FSL MCFLIRT. If more than 3 consecutive images had more than 0.5 millimeters or 0.5 radians of motion, there had to be at least 7 consecutive low-motion volumes following the last high-motion volume in order for those volumes to be included in the analysis. Additionally, each subrun had to have at least 24 volumes after accounting for motion and sleep TRs. Functional data were skull-stripped (FSL BET2), intensity normalized, and spatially smoothed with a 3mm FWHM Gaussian kernel (FSL SUSAN).

### Data registration

Due to the lack of an anatomical image from most subjects, we registered all functional data to a representative functional image collected with the same coil and acquisition parameters (referred to as the functional template image). As there is no standard approach to the use of common space in infant MRI research, we elected to use a template image collected with the same acquisition parameters as each subject. This image was originally collected by and reported in Kosakowski et al. (2022). The full procedure for registering each subrun to functional template space was as follows. First, the middle image of each subrun was extracted and used as an example image for registration. If the middle image was corrupted by motion or distortion, a better image was selected to be the example image. Second, the middle image from each subrun was registered to a target image for each subject using a 6 degree of freedom rigid-body transform. The target image for each subject was defined as the middle image of the first subrun of the first visit. Third, the target image for each subject was registered to the functional template image using a 12 degree of freedom affine transform. Fourth, and finally, the subrun-to-target and target-to-template registrations were concatenated so that each subrun was individually registered to template space.

### Parcels / Search Spaces

Group constrained parcels were acquired from previous adult research to localize areas of selectivity (Julian et al., 2012). Scene-selective parcels included OPA, PPA, and MPA, while the object preferring parcel was the LOC. All parcels included separate left and right hemisphere parcels. Given the difficulty of obtaining reliable registrations with infant functional data, we dilated the OPA parcel to increase the chances of capturing peak voxels in each region. The final parcels are available on OSF.

### Subject-level Beta and Contrast Maps

Functional data were analyzed with a whole brain voxel-wise general linear model (GLM) using the FSL software. The GLM included 4 condition regressors (scenes, objects, faces, and scrambled scenes), 6 motion regressors, a linear trend regressor, and 5 PCA noise regressors. Condition regressors were defined as a boxcar function for the duration of the stimulus presentation (18s blocks). Infant inattention or sleep was accounted for using a single impulse nuisance (‘sleep’) regressor. The sleep regressor was defined as a boxcar function with a 1 for each TR where the infant was not looking at the stimuli, and the corresponding TR for all condition regressors was set to 0. Boxcar condition and sleep regressors were convolved with an infant hemodynamic response function (HRF) that is characterized by a longer time to peak and deeper undershoot compared to the standard adult HRF (Arichi et al., 2012). Where required for ssfROI analyses, data and regressors were concatenated across subruns, run regressors were added to account for differences between runs, and beta values were computed for each condition in a whole-brain voxel-wise GLM. PCA noise regressors were computed using a method similar to GLMDenoise (Kay et al., 2013) as defined in Deen et al. (2017). Specifically, we first defined a “noise pool” of voxels whose activation was unrelated to the experimental paradigm, based on the results of an initial GLM including condition and motion impulse regressors, and PCA was then run on the timecourses from these noise pool voxels, in order to extract time courses for the top 5 principal components, which were then included as nuisance regressors in the GLM. Noise pool voxels were defined as the bottom 1% of voxels showing the weakest activation to the experimental conditions, calculated as the sum of the z-scored betas for the experimental conditions.

### Group random effects analysis

To test whether there was systematic overlap between areas of activation across infants, we conducted group random effects analyses using the full sample of infants from the ssfROI analysis (N=32 infants). For each session, we generated a single activation map using the data from all available subruns, and for subjects with multiple sessions, we averaged these maps across sessions, yielding a single map per subject. A nonparametric one-sample t-test was performed across subjects using FSL’s randomise (5000 permutations, TFCE correction) (Winkler et al., 2014). For visualization and reporting purposes, voxelwise statistical maps for the contrast of scenes > all control conditions were transformed to infant template space (see data registration method above).

### Statistical analyses

Statistical significance was assessed using linear mixed effects models. All models included factors for age (in months, normalized) and head motion (proportion of scrubbed volumes across all subruns, per infant), as well as a random effect of subject. The fixed effect of age and random effect of subject allowed us to account for infants who produced usable datasets at two different timepoints. For analyses including data from more than one region, models included a factor for region. For analyses of the full dataset, models included a factor for experiment. Following our preregistered analysis plan, p values were assessed with one-tailed tests where a specific direction of effect was predicted (e.g., greater responses to scenes than control stimuli; stronger responses in older than younger infants; stronger responses for higher rectilinearity stimuli). Exploratory analyses testing for effects the same direction as the preregistered analyses were also tested using one-tailed tests.

## Supplemental Materials

### Methods for estimating scene experience

How much experience do infants gain in the first two to nine months of life? Using the EMA data, we estimated the amount of experience per day for any given age by multiplying the observed proportions of ego-motion experience responses (i.e., in Main Figure 1A-D) by estimates of the amount of time spent awake per day at each age. Based on prior work and established recommendations for infant sleep (Iglowsetin et al., 2003; Paruthi et al., 2016), we estimated that infant wakefulness increases gradually from 7.5 hrs per day at birth to 10 hrs per day at 9 months. To calculate overall experience with ego-motion through the environment, we multiplied the vector of time spent awake per day by the vector of proportion “yes” ego-motion responses (as a function of age) in Figure 1A. To account for the fact that ego-motion events may differ in length, as a function of age, we further multiplied these vectors by the vector of proportion “yes” responses to ego-motion duration greater than 10 minutes in Figure 1B. Finally, to estimate forward-facing ego-motion experience, we additionally multiplied by the vector of proportion “yes” forward ego-motion responses in Figure 1C. The resultant vectors were interpolated to provide one value per day of life, and cumulative experience was then calculated as the sum of the final vector. Taking this approach, we estimate that infants ages 2 to 9 months old (the age range of our infant fMRI sample) have approximately 146-1427 hours of ego-motion experience, and 62-1223 total hours of forward-facing ego-motion experience (i.e., as depicted in our stimuli). A 4.88 month old (the median age of the full sample) is estimated to have 522 hours of ego-motion experience, and 338 hours of forward facing ego-motion experience. Notably, these figures should be considered an upper bound for potential visual scene experience, given that many aspects of visual perception are quite limited in the first few months, including visual acuity (impacting high spatial frequency and rectilinearity experience) and peripheral vision (Daw and Daw, 2006; Atkinson, 2002).

### Additional analyses of scene selectivity

To further test infant scene selectivity, here we report comparisons of responses to scenes versus each control condition, separately in each region. Note that estimates of the responses to individual control conditions are based on less data, since each control condition was sampled 3x less often than scenes in the preregistered dataset. Note also that the set of conditions differed between the two samples; both included scenes, faces, and objects, while only the preregistered sample included scrambled scenes and only the Kosakowski et al. (2022) sample included bodies.

Results are summarized in Figure 3. Beginning with the preregistered dataset, for OPA, we found greater responses to scenes than objects (β = -15.39, t_(59.46)_ = -1.97, p = 0.03) and faces (β = -23.87, t_(59.46)_ = -3.05, p = 0.002), although the comparison to scrambled scenes fell short of significance (β = -11.26, t_(59.46)_ = -1.44, p = 0.08). For PPA, we found greater responses to scenes than faces (β = -6.91, t_(62.93)_ = -2.50, p = 0.007) and scrambled scenes (β = -7.31, t_(62.93)_ = -2.65, p = 0.005), but not objects (β = 0.54, t_(62.93)_ = 0.20, p = 0.84; two-tailed test reported, rather than preregistered one-tailed test, since the response to objects was numerically larger). For MPA, we found greater responses to scenes than objects (β = -9.556, t_(57.80)_ = -2.395, p = 0.01) and scrambled scenes (β = -10.198, t_(57.80)_ = -2.556, p = 0.007), while the comparison to faces fell short of significance (β = -5.405, t_(57.80)_ = -1.355, p = 0.09). Next, in the full sample of infants, for OPA, we found greater responses to scenes than faces (β = -22.99, t_(105.23)_ = -3.38, p < 0.001) and bodies (β = -18.26, t_(105.23)_ = -1.93, p = 0.03), with a trend that did not reach significance relative to objects (β = -9.88, t_(105.23)_ = -1.45, p = 0.07), and no effect relative to scrambled scenes (β = -9.13, t_(105.23)_ = -1.11, p = 0.14). For PPA, we found greater responses to scenes than faces (β = -8.57, t_(105.23)_ = -4.00, p < 0.001), scrambled scenes (β = -8.97, t_(105.23)_ = -3.44, p < 0.001), and bodies (β = -5.08, t_(105.23)_ = -1.70, p = 0.046), with a trend that did not reach significance relative to objects (β = -2.77, t_(105.23)_ = -1.29, p = 0.099). Finally, for MPA, we found greater responses to scenes than objects (β = -8.54, t_(101.74)_ = -2.87, p = 0.003), faces (β = -6.06, t_(101.74)_ = -2.03, p = 0.02), and scrambled scenes (β = -10.08, t_(101.74)_ = -2.78, p = 0.003), but not bodies (β = -5.02, t_(101.74)_ = -1.21, p = 0.11).

Lastly, we found no evidence of reliable differences in the full functional profiles of the three scene regions; an exploratory ANOVA failed to find a region x condition interaction in either the preregistered (F_(6,218.01)_ = 1.05, p = 0.39) or full sample (F_(8,382.90)_ = 1.04, p = 0.40).

### Additional analyses of low-level visual features

To further test whether responses in infant scene regions were better explained by scene selectivity vs. low-level features, we conducted analyses in individual scene regions, noting that the pattern of responses across conditions did not differ between the regions in the analyses above. To do this, we used linear mixed effect models to test whether a categorical factor for scene selectivity explains significant variance in responses even when accounting for variance explained by each low-level control feature). For *<u>OPA</u>*, the effect of category was significant after accounting for each of the four low-level feature models, in both the full and preregistered datasets (Preregistered: all β’s > 5.84, all t’s_(107.29)_ > 1.76, all p’s < 0.04; Full: all β’s > 7.16, all t’s_(60.44)_ > 2.16, all p’s < 0.02). No low-level visual feature explaining significant variance (Preregistered: all β’s < 4.44, all t’s_(60.44)_ < 1.55, all p’s > 0.06, Full: all β’s < 3.08, all t’s_(107.29)_ < 1.00, all p’s > 0.33). For *<u>PPA</u>*, the effect of category held accounting for high spatial frequency, overall motion energy, and peripheral model energy in both datasets (Preregistered: all β’s > 2.45, all t’s_(63.94)_ > 1.98, all p’s < 0.03; Full: all β’s > 3.18, all t’s_(109.19)_ > 2.96, all p’s < 0.002). None of these low-level visual features explained significant variance (Preregistered: all β’s < -0.63, all t’s_(63.94)_ < -0.59, all p’s > 0.17; Full: all β’s < 0.12, all t’s_(109.19)_ < 0.12, all p’s > 0.61). However, there was a significant effect of rectilinearity in both datasets (Preregistered: β = 3.35, t_(63.93)_ = 3.17, p = 0.001; Full: β = 2.57, t_(109.19)_ = 2.85, p = 0.003), which explained the category effect in the preregistered dataset (β = 0.881, t_(63.93)_ = 0.67, p = 0.25), but not the full dataset (Full: β = 2.57, t_(109.19)_ = 2.85, p = 0.003). For *<u>MPA</u>*, the effect of category was significant after accounting for each of the four low-level feature models (Preregistered: all β’s > 4.48, all t’s_(58.81)_ > 2.62, all p’s < 0.006; Full: all β’s > 3.91, all t’s_(103.63)_ > 3.13, all p’s < 0.002), with no low-level visual feature explaining significant variance (Preregistered: all β’s < -1.03, all t’s_(58.81)_ < -0.66, all p’s > 0.13; Full: all β’s < -0.67, all t’s_(103.63)_ < -0.54, all p’s > 0.11). Taken together, at the level of individual regions, we again found no reliable evidence that scene regions are better explained by low-level visual features than scene selectivity.

**Supplemental Figure 1.**
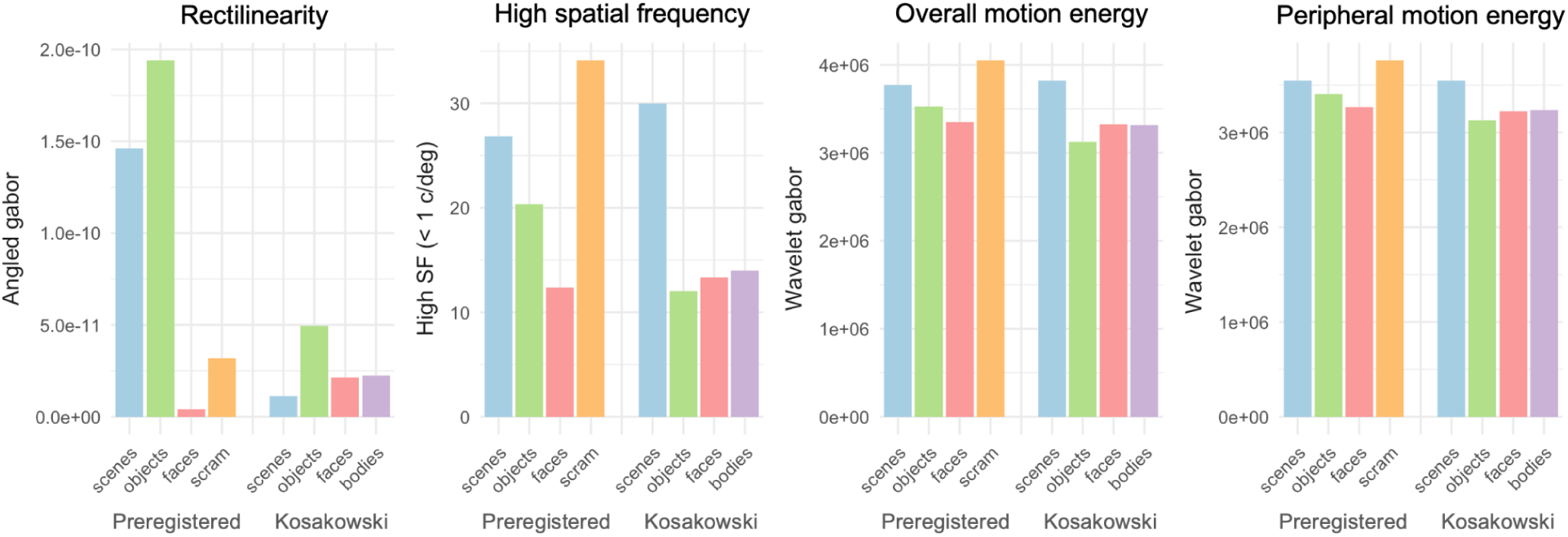
*Average low-level visual feature content per stimulus category*. Rectilinearity, high spatial frequency, and motion energy (overall and in the periphery of the video frame) were estimated for each frame/stimulus and averaged within categories (see Methods). Each plot shows mean feature values for the stimuli in the preregistered dataset (left four bars in each plot) and in Kosakowski et al. (2022) (right four bars in each plot).

### Additional analyses of age-related change

Here we report two further analyses of age related change. First, to provide a broader test of age-related change for any individual condition, we tested for an interaction of age and each of the conditions in each region. For the preregistered dataset, we found no evidence of age-related change for any individual condition in OPA (all β’s < 13.97, all t’s < 1.58, all p’s > 0.13), PPA (all β’s < 2.74, all t’s < 1.20, all p’s > 0.24) or MPA (all β’s < 6.87, all t’s < 1.47, all p’s > 0.15). Likewise, with the full dataset, we found no evidence of age-related change for any individual condition in OPA (all β’s < 9.97, all t’s < 1.51, all p’s > 0.13), PPA (all β’s < 2.73, all t’s < 1.20, all p’s > 0.24) or MPA (all β’s < 7.14, all t’s < 1.69, all p’s > 0.09).

Second, we report tests of scene selectivity in the younger and older (median split) age groups, separately for each scene region. Age data for the median split of the full sample is presented in the main text. For the preregistered dataset, the younger sample included 11 sessions from 10 infants (mean age = 3.60 months, range = 2.07-5.33 months). As with the full sample, an exploratory omnibus analysis in the preregistered sample revealed a significantly greater response to scenes than control conditions (β = 4.28, t_(117.12)_ = 1.69, p = 0.046).

We also performed these analyses on individual regions, noting the limited power of these analyses, and lack of evidence for reliable differences between regions. For younger infants in the preregistered dataset, OPA showed significantly greater responses to scenes than control stimuli in younger infants (β = 8.41, t_(29.92)_ = 2.24, p = 0.02), with a similar, non-significant trend in older infants (β = 8.43, t_(30.52)_ = 1.58, p = 0.06). PPA showed significantly greater responses to scenes than control stimuli in older infants (β = 3.44, t_(29.83)_ = 1.90, p = 0.03) but not younger infants (β = 1.22, t_(32.69)_ = 0.77, p = 0.22). MPA showed significantly greater responses to scenes than control stimuli in older infants (β = 5.26, t_(29.69)_ = 2.22, p = 0.02), with a similar, non-significant trend in younger infants (β = 3.22, t_(31.89)_ = 1.63, p = 0.06). No region showed a significant age group x scene selectivity interaction (all β’s < 1.11, all t’s < 0.93, all p’s > 0.35). For exploratory analyses of the full dataset, OPA showed significantly greater responses to scenes than control stimuli in older infants (β = 9.50, t_(53.70)_ = 2.27, p = 0.01), with a similar trend that did not reach significance in younger infants (β = 5.97, t_(55.48)_ = 1.65, p = 0.052). PPA showed significantly greater responses to scenes than control stimuli in both younger (β = 2.54, t_(56.27)_ = 1.99, p = 0.03) and older infants (β = 3.76, t_(51.34)_ = 2.93, p = 0.003). MPA showed significantly greater responses to scenes than control stimuli in older infants (β = 6.15, t_(51.72)_ = 3.44, p < 0.001), but not younger infants (β = 1.63, t_(56.68)_ = 1.07, p = 0.14). We failed to find a significant age group x scene selectivity interaction in MPA (β = 2.26, t_(109.56)_ = 1.88, p = 0.06), OPA (β = 1.77, t_(109.32)_ = 0.64, p = 0.53) or PPA (β = 0.61, t_(109.33)_ = 0.67, p = 0.50). Taken together, these analyses failed to reveal clear evidence of age-related change in cortical scene responses during the first nine months (despite some evidence of trends toward increasing selectivity in MPA). Rather, these data suggest that scene selective responses are relatively stable across the first nine months, and already present as early as 2-5 months old.

## References

Arcaro, M. J., & Livingstone, M. S. (2017). A hierarchical, retinotopic proto-organization of the primate visual system at birth. Elife, 6, e26196.

Arcaro, M. J., & Livingstone, M. S. (2021). On the relationship between maps and domains in inferotemporal cortex. Nature Reviews Neuroscience, 22(9), 573–583.

Arichi, T., Fagiolo, G., Varela, M., Melendez-Calderon, A., Allievi, A., Merchant, N., … & Edwards, A. D. (2012). Development of BOLD signal hemodynamic responses in the human brain. Neuroimage, 63(2), 663–673.

Atkinson, Janette, ’Newborn vision’, The Developing Visual Brain, Oxford Psychology Series (Oxford, 2002; online edn, Oxford Academic, 1 Jan. 2008), 10.1093/acprof:oso/9780198525998.003.0004.

Bryan, P. B., Julian, J. B., & Epstein, R. A. (2016). Rectilinear edge selectivity is insufficient to explain the category selectivity of the parahippocampal place area. Frontiers in human neuroscience, 10, 137.

Cabral, L., Zubiaurre-Elorza, L., Wild, C. J., Linke, A., & Cusack, R. (2022). Anatomical correlates of category-selective visual regions have distinctive signatures of connectivity in neonates. Developmental Cognitive Neuroscience, 58, 101179.

Chen, E. M.*, Kamps, F. S.*, & Saxe, R. R. (2024). An open dataset of functional MRI responses to egocentric navigation through natural scenes. 7th Annual Conference on Cognitive Computational Neuroscience. Cambridge, Massachusetts, USA.

Daw, N. W., & Daw, N. W. (2006). Visual development (Vol. 14). New York: Springer.

Deen, B., Richardson, H., Dilks, D. D., Takahashi, A., Keil, B., Wald, L. L., … & Saxe, R. (2017). Organization of high-level visual cortex in human infants. Nature communications, 8(1), 13995.

Dilks, D. D., Julian, J. B., Paunov, A. M., & Kanwisher, N. (2013). The occipital place area is causally and selectively involved in scene perception. Journal of neuroscience, 33(4), 1331–1336.

Dilks, D. D., Jung, Y., & Kamps, F. S. (2023). The development of human cortical scene processing. Current directions in psychological science, 32(6), 479–486.

Dilks, D. D., Kamps, F. S., & Persichetti, A. S. (2022). Three cortical scene systems and their development. Trends in cognitive sciences, 26(2), 117–127.

Epstein, R. A., & Baker, C. I. (2019). Scene perception in the human brain. Annual review of vision science, 5(1), 373–397.

Epstein, R., & Kanwisher, N. (1998). A cortical representation of the local visual environment. Nature, 392(6676), 598–601.

Ghotra, A., Kosakowski, H. L., Takahashi, A., Etzel, R., May, M. W., Scholz, A., … & Keil, B. (2021). A size-adaptive 32-channel array coil for awake infant neuroimaging at 3 Tesla MRI. Magnetic Resonance in Medicine, 86(3), 1773–1785.

Gomez, J., Natu, V., Jeska, B., Barnett, M., & Grill-Spector, K. (2018). Development differentially sculpts receptive fields across early and high-level human visual cortex. Nature communications, 9(1), 788.

Gibson, E. J., & Walk, R. D. (1960). The “visual cliff”. Scientific American, 202(4), 64–71.

Gilmore, R. O., Raudies, F., & Jayaraman, S. (2015, August). What accounts for developmental shifts in optic flow sensitivity?. In 2015 Joint IEEE International Conference on Development and Learning and Epigenetic Robotics (ICDL-EpiRob) (pp. 19–25). IEEE.

Golarai, G., Ghahremani, D. G., Whitfield-Gabrieli, S., Reiss, A., Eberhardt, J. L., Gabrieli, J. D., & Grill-Spector, K. (2007). Differential development of high-level visual cortex correlates with category-specific recognition memory. Nature neuroscience, 10(4), 512–522.

Hasson, U., Harel, M., Levy, I., & Malach, R. (2003). Large-scale mirror-symmetry organization of human occipito-temporal object areas. Neuron, 37(6), 1027–1041.

He, C., Peelen, M. V., Han, Z., Lin, N., Caramazza, A., & Bi, Y. (2013). Selectivity for large nonmanipulable objects in scene-selective visual cortex does not require visual experience. Neuroimage, 79, 1–9.

Held, R., & Hein, A. (1963). Movement-produced stimulation in the development of visually guided behavior. Journal of comparative and physiological psychology, 56(5), 872.

Hermer, L., & Spelke, E. (1996). Modularity and development: The case of spatial reorientation. Cognition, 61(3), 195–232.

Jiang, P., Tokariev, M., Aronen, E. T., Salonen, O., Ma, Y., Vuontela, V., & Carlson, S. (2014). Responsiveness and functional connectivity of the scene-sensitive retrosplenial complex in 7–11-year-old children. Brain and cognition, 92, 61–72.

Julian, J. B., Fedorenko, E., Webster, J., & Kanwisher, N. (2012). An algorithmic method for functionally defining regions of interest in the ventral visual pathway. Neuroimage, 60(4), 2357–2364.

Jung, Y., Hsu, D., & Dilks, D. D. (2024). “Walking selectivity” in the occipital place area in 8-year-olds, not 5-year-olds. Cerebral Cortex, 34(3), bhae101.

Kamps, F. S., Hendrix, C. L., Brennan, P. A., & Dilks, D. D. (2020). Connectivity at the origins of domain specificity in the cortical face and place networks. Proceedings of the National Academy of Sciences, 117(11), 6163–6169.

Kamps, F. S., Lall, V., & Dilks, D. D. (2016). The occipital place area represents first-person perspective motion information through scenes. Cortex, 83, 17–26.

Kamps, F. S., Pincus, J. E., Radwan, S. F., Wahab, S., & Dilks, D. D. (2020). Late development of navigationally relevant motion processing in the occipital place area. Current Biology, 30(3), 544–550.

Kamps, F. S.*, Rennert, R. J.*, Radwan, S. F., Wahab, S., Pincus, J. E., & Dilks, D. D. (2023). Dissociable cognitive systems for recognizing places and navigating through them: Developmental and neuropsychological evidence. Journal of Neuroscience, 43(36), 6320–6329.

Kamps, F. S., Richardson, H., Murty, N. A. R., Kanwisher, N., & Saxe, R. (2022). Using child-friendly movie stimuli to study the development of face, place, and object regions from age 3 to 12 years. Human Brain Mapping, 43(9), 2782–2800.

Kay, K. N., Rokem, A., Winawer, J., Dougherty, R. F., & Wandell, B. A. (2013). GLMdenoise: a fast, automated technique for denoising task-based fMRI data. Frontiers in neuroscience, 7, 247.

Kosakowski, H. L., Cohen, M. A., Takahashi, A., Keil, B., Kanwisher, N., & Saxe, R. (2022). Selective responses to faces, scenes, and bodies in the ventral visual pathway of infants. Current Biology, 32(2), 265–274.

Kosakowski, H. L., Cohen, M. A., Herrera, L., Nichoson, I., Kanwisher, N., & Saxe, R. (2024). Cortical face-selective responses emerge early in human infancy. eneuro, 11(7).

Kravitz, D. J., Peng, C. S., & Baker, C. I. (2011). Real-world scene representations in high-level visual cortex: it’s the spaces more than the places. Journal of Neuroscience, 31(20), 7322–7333.

Kretch, K. S., Franchak, J. M., & Adolph, K. E. (2014). Crawling and walking infants see the world differently. Child development, 85(4), 1503–1518.

Levy, I., Hasson, U., Harel, M., & Malach, R. (2004). Functional analysis of the periphery effect in human building related areas. Human brain mapping, 22(1), 15–26.

Levy, I., Hasson, U., Avidan, G., Hendler, T., & Malach, R. (2001). Center–periphery organization of human object areas. Nature neuroscience, 4(5), 533–539.

Li, J., Osher, D. E., Hansen, H. A., & Saygin, Z. M. (2020). Innate connectivity patterns drive the development of the visual word form area. Scientific reports, 10(1), 18039.

Maguire, E. (2001). The retrosplenial contribution to human navigation: a review of lesion and neuroimaging findings. Scandinavian journal of psychology, 42(3), 225–238.

Nasr, S., Echavarria, C. E., & Tootell, R. B. (2014). Thinking outside the box: rectilinear shapes selectively activate scene-selective cortex. Journal of Neuroscience, 34(20), 6721–6735.

Nishimoto, S., Vu, A. T., Naselaris, T., Benjamini, Y., Yu, B., & Gallant, J. L. (2011). Reconstructing visual experiences from brain activity evoked by natural movies. Current biology, 21(19), 1641–1646.

Orhan, A. E., & Lake, B. M. (2024). Learning high-level visual representations from a child’s perspective without strong inductive biases. Nature Machine Intelligence, 6(3), 271–283.

Park, S., Brady, T. F., Greene, M. R., & Oliva, A. (2011). Disentangling scene content from spatial boundary: complementary roles for the parahippocampal place area and lateral occipital complex in representing real-world scenes. Journal of Neuroscience, 31(4), 1333–1340.

Park, S., Konkle, T., & Oliva, A. (2015). Parametric coding of the size and clutter of natural scenes in the human brain. Cerebral cortex, 25(7), 1792–1805.

Paruthi, S., Brooks, L. J., D’Ambrosio, C., Hall, W. A., Kotagal, S., Lloyd, R. M., … & Wise, M. S. (2016). Recommended amount of sleep for pediatric populations: a consensus statement of the American Academy of Sleep Medicine. Journal of clinical sleep medicine, 12(6), 785–786.

Petroff, Z. J., Jayaraman, S., Smith, L. B., Candy, T. R., & Bonnen, K. (2025). The world through infant eyes: Evidence for the early emergence of the cardinal orientation bias. Proceedings of the National Academy of Sciences, 122(16), e2421277122.

Pitcher, D., Dilks, D. D., Saxe, R. R., Triantafyllou, C., & Kanwisher, N. (2011). Differential selectivity for dynamic versus static information in face-selective cortical regions. Neuroimage, 56(4), 2356–2363.

Powell, L. J., Deen, B., & Saxe, R. (2018). Using individual functional channels of interest to study cortical development with fNIRS. Developmental science, 21(4), e12595.

Powell, L. J., Kosakowski, H. L., & Saxe, R. (2018). Social origins of cortical face areas. Trends in cognitive sciences, 22(9), 752–763.

Rajimehr, R., Devaney, K. J., Bilenko, N. Y., Young, J. C., & Tootell, R. B. (2011). The “parahippocampal place area” responds preferentially to high spatial frequencies in humans and monkeys. PLoS biology, 9(4), e1000608.

Raudies, F., Gilmore, R. O., Kretch, K. S., Franchak, J. M., & Adolph, K. E. (2012, November). Understanding the development of motion processing by characterizing optic flow experienced by infants and their mothers. In 2012 IEEE International Conference on Development and Learning and Epigenetic Robotics (ICDL) (pp. 1–6). IEEE.

Raudies, F., & Gilmore, R. O. (2014). Visual motion priors differ for infants and mothers. Neural computation, 26(11), 2652–2668.

Raz, G., & Saxe, R. (2020). Learning in infancy is active, endogenously motivated, and depends on the prefrontal cortices. Annual Review of Developmental Psychology, 2(1), 247–268.

Scherf, K. S., Behrmann, M., Humphreys, K., & Luna, B. (2007). Visual category-selectivity for faces, places and objects emerges along different developmental trajectories. Developmental science, 10(4), F15–F30.

Scherf, K. S., Luna, B., Avidan, G., & Behrmann, M. (2011). “What” precedes “which”: developmental neural tuning in face-and place-related cortex. Cerebral cortex, 21(9), 1963–1980.

Silson, E. H., Steel, A. D., & Baker, C. I. (2016). Scene-selectivity and retinotopy in medial parietal cortex. Frontiers in human neuroscience, 10, 412.

Walther, D. B., Chai, B., Caddigan, E., Beck, D. M., & Fei-Fei, L. (2011). Simple line drawings suffice for functional MRI decoding of natural scene categories. Proceedings of the National Academy of Sciences, 108(23), 9661–9666.

*Winkler* *AM*, *Ridgway* *GR*, *Webster* *MA*, *Smith* *SM*, *Nichols* *TE*. Permutation inference for the general linear model. NeuroImage, 2014;92:381-397. (Open Access)

Wolbers, T., Klatzky, R. L., Loomis, J. M., Wutte, M. G., & Giudice, N. A. (2011). Modality-independent coding of spatial layout in the human brain. Current Biology, 21(11), 984–989.

